# AINN-Express: A Leakage-Aware, Sequence-Only Predictor of VHH Antibody Expression Built on the AINN-P1 Protein Foundation Model

**DOI:** 10.64898/2026.07.21.739256

**Authors:** Roger Wang, Kevin Jin, Lurong Pan

## Abstract

Expression — whether an antibody can be produced at usable yield — is one of the earliest and most expensive filters in therapeutic discovery. We present **AINN-Express**, a sequence-only predictor of VHH single-domain antibody (nanobody) expression built on **AINN-P1**, Ainnocence’s protein foundation model. AINN-Express encodes a VHH with a frozen AINN-P1 encoder and scores it with a lightweight gradient-boosted classifier: it takes only an amino-acid sequence, returns an expression probability, and needs no structure and no per-task model training. Under a leakage-safe, leave-program-out evaluation, AINN-Express reaches ROC-AUC 0.87 within known antibody programs and 0.81 on entirely new programs — well above the majority baseline — making it a practical tool for prioritizing candidates before wet-lab work. We further justify the encoder choice with a controlled, leakage-aware benchmark against general-purpose protein language models: on this task, AINN-P1 (167 M parameters) generalizes to unseen programs far better than a general-purpose ESM2 (650 M) — 0.81 versus 0.68 new-program ROC-AUC — and matches a domain-finetuned ESM2 (0.83) with no task-specific finetuning, at roughly one-quarter of the parameters. The gap is invisible under a random split, where all encoders score ∼0.88; only leave-program-out evaluation reveals that general-purpose embeddings largely encode *program identity* rather than transferable determinants of expression. Purpose-built representation quality, not parameter count, is what makes AINN-Express generalize.

## 1. Introduction

Producing an antibody at usable yield is a prerequisite for almost everything downstream in discovery, yet expression is typically discovered only at the bench. A reliable sequence-only predictor lets teams rank and triage candidates in silico, spending scarce wet-lab capacity on the most promising ones. Protein language models (pLMs) make this practical: embeddings from a model pretrained on large sequence corpora transfer to many downstream properties without structural supervision (Lin et al., 2023; Rives et al., 2021).

Two questions then follow. The first is practical: can a simple, deployable pipeline predict VHH expression from sequence alone accurately enough to be useful for triage? The second is about representation: as purpose-built foundation models emerge alongside ever-larger general-purpose ones, does a domain-oriented encoder produce more transferable representations for a narrow domain such as antibody engineering than a larger general-purpose one? This manuscript answers both, and shows they are connected — the tool works because the encoder choice is right, and the difference only becomes visible under leakage-aware evaluation.

We introduce AINN-Express, a VHH expression predictor built on AINN-P1, and pair it with the encoder benchmark that motivates its design. Our contributions are:

- **AINN-Express**: a sequence-only VHH expression predictor — frozen AINN-P1 embeddings and a gradient-boosted classifier — designed for candidate triage, with a leakage-safe evaluation showing ROC-AUC 0.87 within known programs and 0.81 on unseen programs, far above the majority baseline.
- A **leakage-aware encoder benchmark** establishing that AINN-P1 embeddings generalize markedly better than general-purpose ESM2 on this task (+0.13 new-program AUC) at roughly one-quarter of the parameters, and match a domain-finetuned ESM2 without any finetuning.
- A mechanistic account of **why** — general-purpose embeddings encode program identity, a signal random splits reward and program-grouped splits expose — and a clear **intended-use framing**: prioritization with a human and experiment in the loop, not an automated hard filter.

## 2. AINN-Express: a sequence-only expression predictor

### 2.1 Pipeline

An overview of AINN-Express is shown in Figure 1. A VHH sequence is encoded by a frozen AINN-P1 foundation model into a single fixed-dimensional embedding, which a lightweight gradient-boosted classifier maps to an expression probability. The design takes only an amino-acid sequence, requires no structure, and trains no part of the foundation model. Freezing the encoder makes the model fast, reproducible, and data-efficient, and requires no GPU training per task; once embeddings are computed, the classifier trains in seconds on CPU.

**Figure 1.**
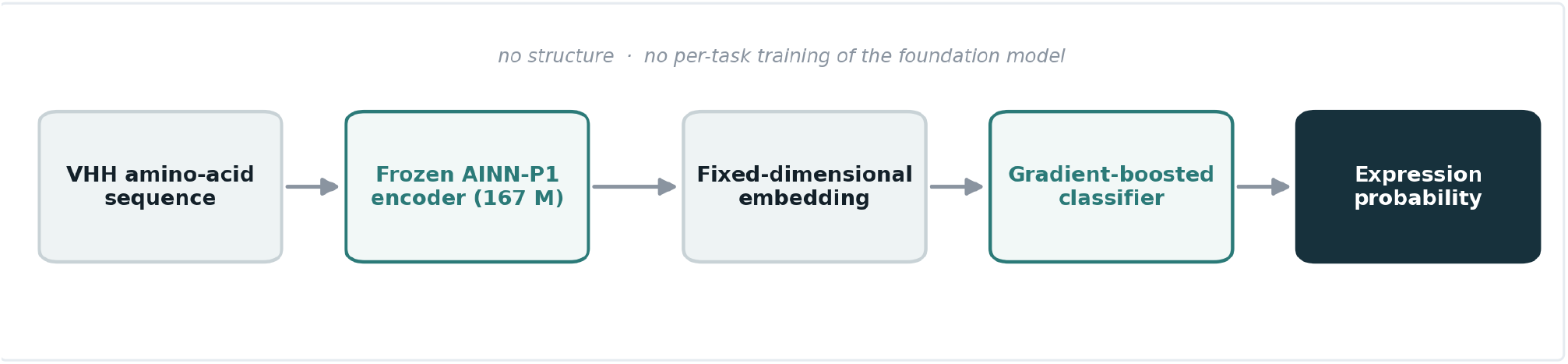
AINN-Express pipeline. A VHH amino-acid sequence is encoded by a frozen AINN-P1 foundation model into a fixed-dimensional embedding, which a lightweight gradient-boosted classifier maps to an expression probability.

### 2.2 Performance

We evaluated AINN-Express under two protocols on a proprietary VHH expression panel (Table 1, Figure 2), reporting standard classification metrics: ROC-AUC, accuracy, F1, precision, recall, and the Matthews correlation coefficient (MCC). Within known antibody programs it reaches ROC-AUC 0.87 (accuracy 0.79); on entirely new programs it holds at ROC-AUC 0.81 (accuracy 0.71) — both far above the majority baseline (AUC 0.50, accuracy 0.54). An AUC of 0.81 means that, given a random expressing and a random non-expressing VHH from an unseen program,the model ranks them correctly 81% of the time — strong enrichment for triage. AUC and MCC are the primary metrics, being threshold-independent and robust to the mild class imbalance; thresholded metrics are reported at a 0.5 cutoff.

**Table 1.** AINN-Express performance on VHH expression. pooled over out-of-fold predictions. The new-program row is the honest generalization estimate; thresholded metrics use a 0.5 cutoff.

| Setting | ROC-AUC | Accuracy | F1 | Precision | Recall | MCC |
| --- | --- | --- | --- | --- | --- | --- |
| Within known programs (random split) | 0.87 | 0.79 | 0.82 | 0.80 | 0.84 | 0.58 |
| New programs (leave-program-out) | 0.81 | 0.71 | 0.74 | 0.71 | 0.78 | 0.40 |
| Majority-class baseline | 0.50 | 0.54 | — | — | — | — |

**Figure 2.**
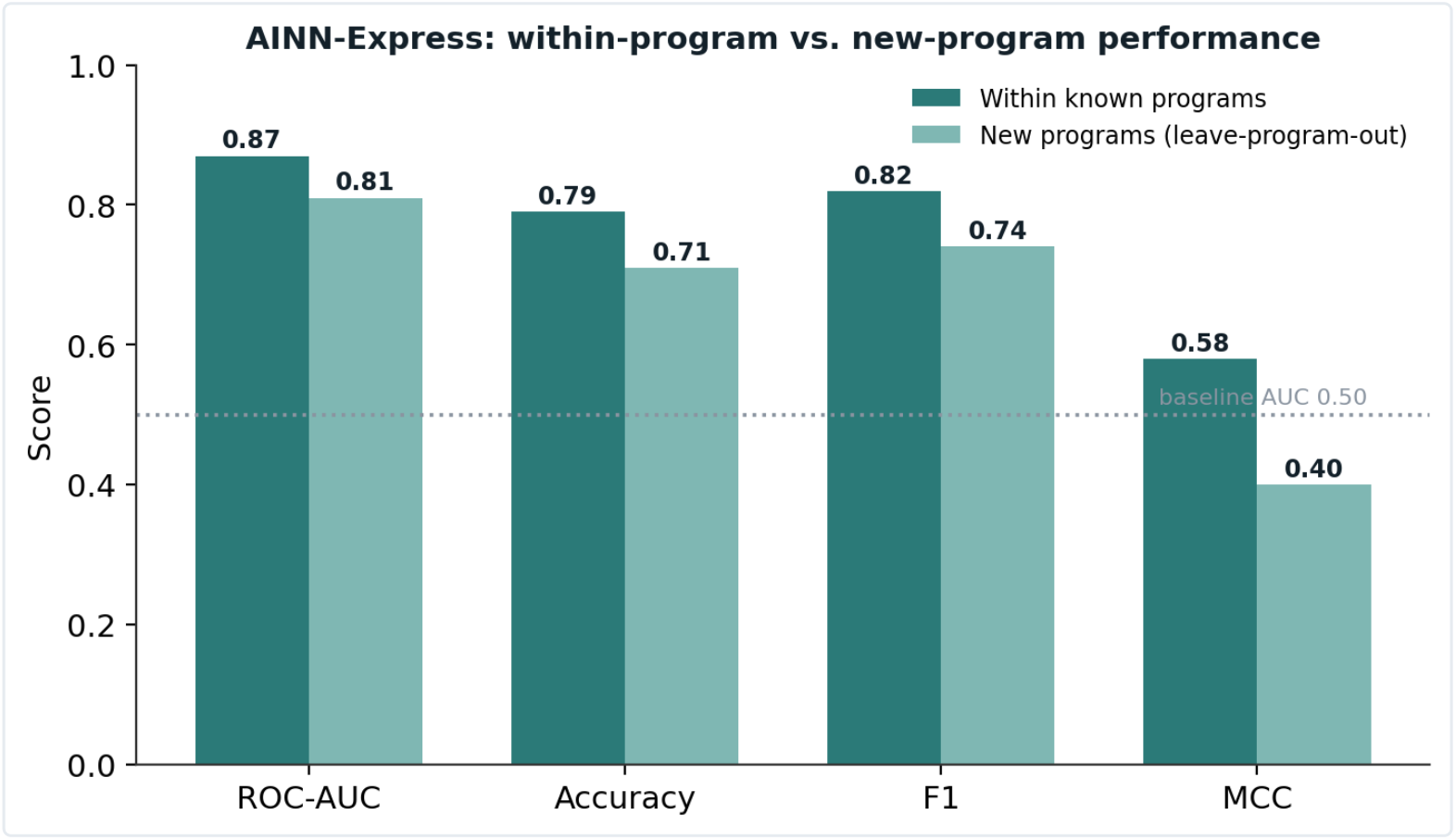
AINN-Express performance within known programs versus on new programs, across ROC-AUC, accuracy, F1, and MCC. All metrics remain well above the majority baseline on unseen programs.

### 2.3 Classifier selection

We selected gradient-boosted trees by comparing a family of lightweight heads on the same AINN-P1 embeddings under leave-program-out cross-validation (Figure 3). Tree ensembles clearly outperform linear heads — the AINN-P1 embeddings are informative but not linearly separable across programs — and gradient boosting is the single strongest choice, which is what AINN-Express deploys.

**Figure 3.**
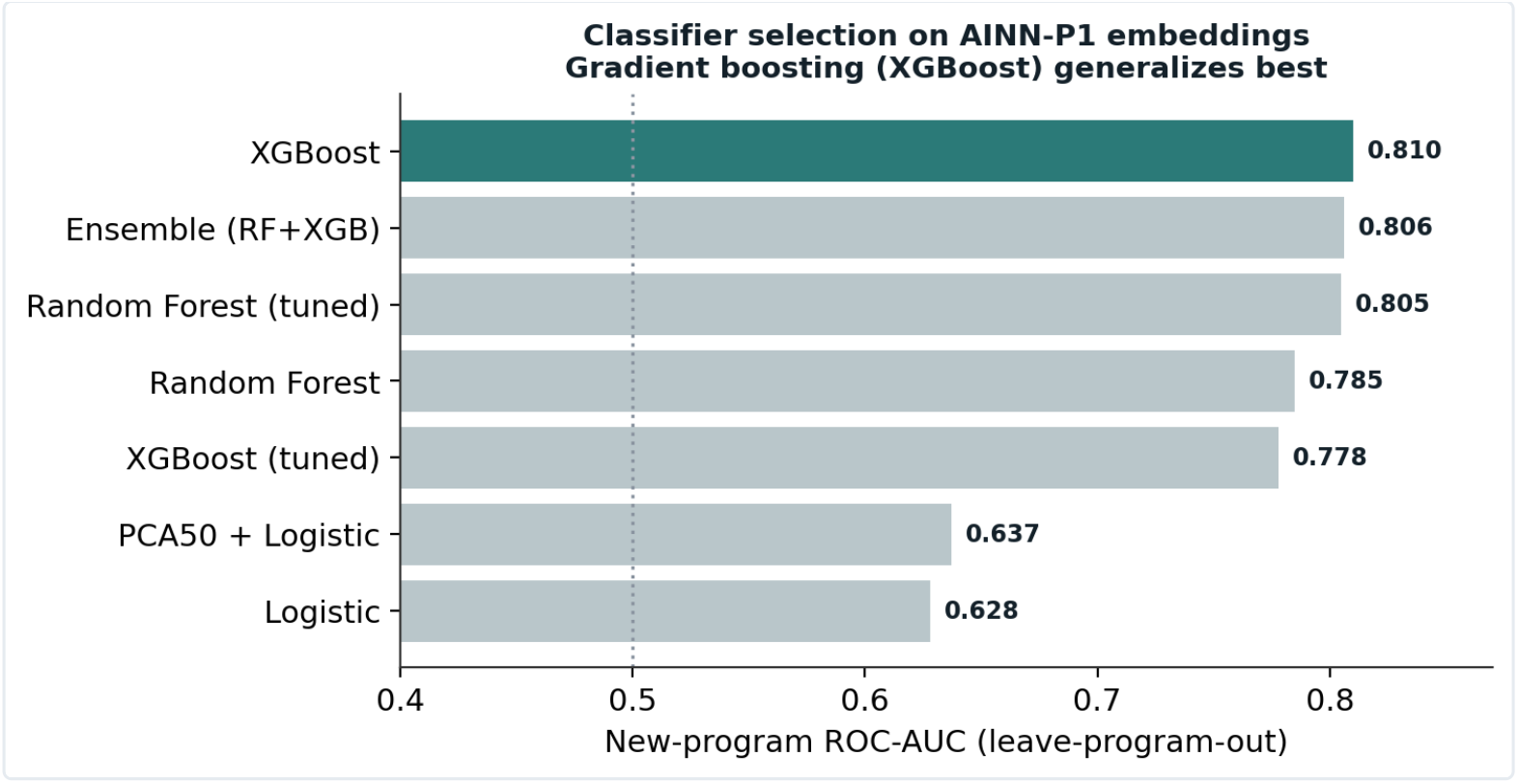
Classifier selection on AINN-P1 embeddings. New-program (leave-program-out) ROC-AUC of candidate classifier heads on frozen AINN-P1 embeddings. Gradient boosting (XGBoost, highlighted) generalizes best and is the head AINN-Express deploys.

## 3. Why AINN-P1? A leakage-aware encoder benchmark

AINN-Express’s generalization to unseen programs rests on the AINN-P1 representation. To justify that choice — and to quantify what it buys — we benchmarked AINN-P1 against two ESM2 baselines under identical downstream treatment: a general-purpose ESM2 (650 M) and a domain-finetuned ESM2 (650 M). Each frozen encoder was paired with the same family of seven lightweight classifiers and scored on identical folds under both a random (within-program) split and a grouped, leave-program-out split. The full 21-configuration table appears as Table 2.

### 3.1 AINN-P1 generalizes to new programs at one-quarter the parameters

On unseen programs, the best AINN-P1 configuration attains ROC-AUC 0.81, well above the best general-purpose ESM2 at 0.68 and comparable to the best domain-finetuned ESM2 at 0.83 (Figure 4), at 167 M parameters — roughly one-quarter of the 650 M in either ESM2 variant. The +0.13 AUC advantage over the general-purpose ESM2 is the central result. Crucially, the same figure exposes why the gap is easy to miss: under a random within-program split all three encoders land within a narrow band around 0.87–0.90, statistically indistinguishable at this panel size. Only the leave-program-out score separates them, and the per-encoder within-minus-new gap makes the leakage explicit — small for AINN-P1 (Δ0.057) and the finetuned ESM2 (Δ0.054), large for the general-purpose ESM2 (Δ0.209).

**Figure 4.**
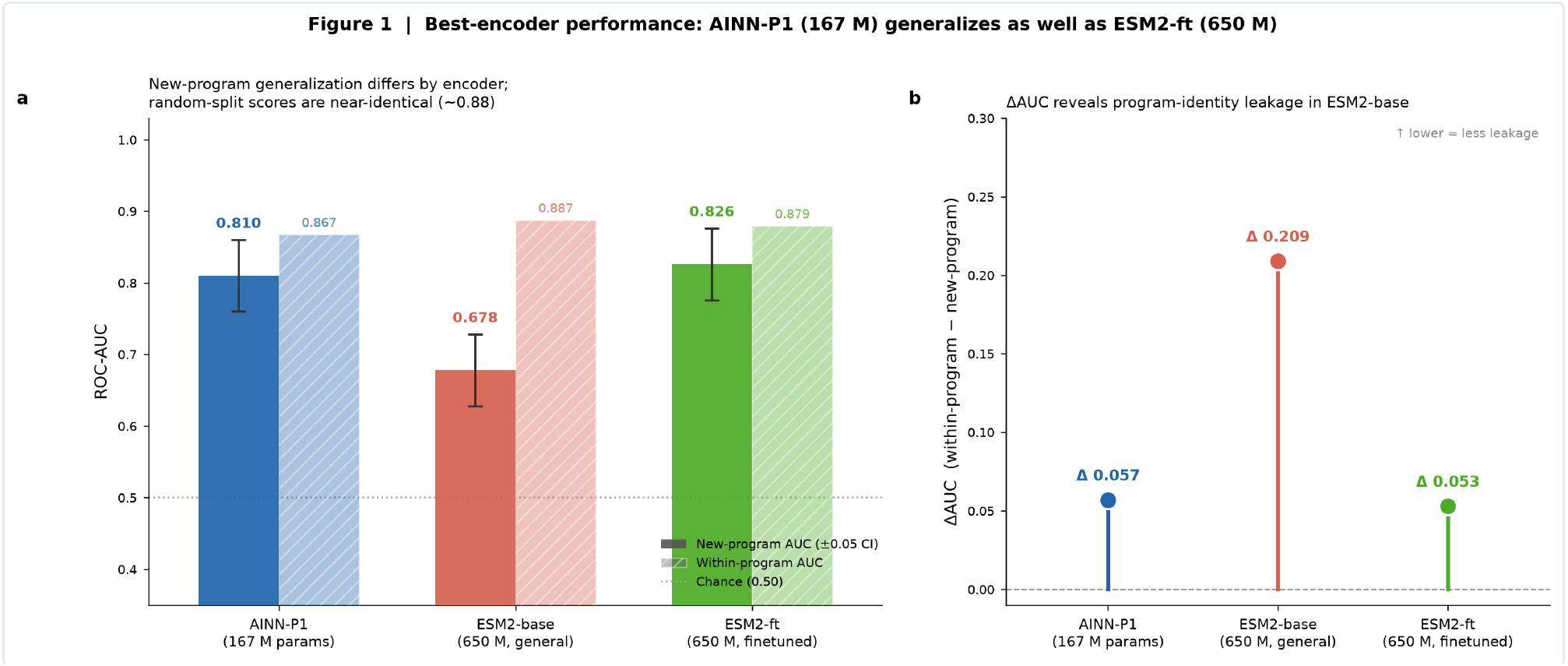
Best-encoder new-program generalization. (A) AINN-P1 (167 M) reaches 0.81 on unseen programs, above the general-purpose ESM2 (0.68) and comparable to the domain-finetuned ESM2 (0.83); random-split scores are near-identical (∼0.88, hatched). (B) The within ™ new-program AUC gap reveals program-identity leakage: small for AINN-P1 (Δ0.057) and ESM2-ft (Δ0.054), large for ESM2-base (Δ0.209). Error bars: ±0.05 CI.

### 3.2 Random-split success does not predict transfer

Plotting each configuration by its within-program AUC against its new-program AUC draws the leakage geometry directly (Figure 5). All three encoders reach ∼0.85–0.90 within programs, so a random-split-only study would rank a 650 M general-purpose model as competitive with a 167 M domain model. The new-program axis — accessible only through leave-program-out evaluation — is where the encoders actually differ: the general-purpose ESM2 falls furthest below the diagonal, its linear heads dropping toward chance on new programs. This is the single most important methodological takeaway: within-program AUC is not a proxy for the quantity that matters in deployment, which is performance on antibody programs the model has never seen.

**Figure 5.**
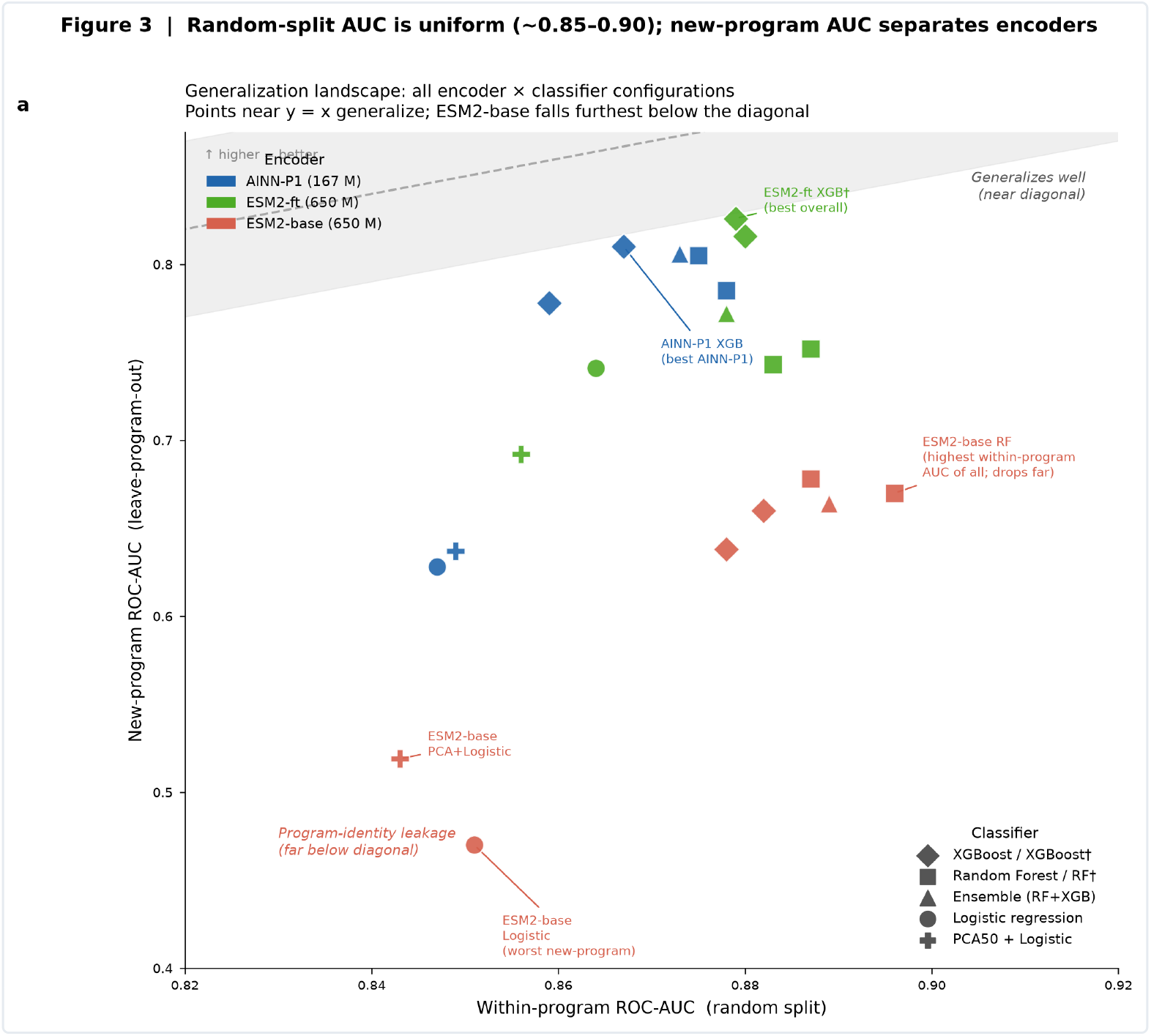
Generalization landscape. Within-program ROC-AUC (x) versus new-program ROC-AUC (y) for all 21 configurations, colored by encoder and shaped by classifier family. Points near y = x generalize; points far below leak on new programs. The general-purpose ESM2 falls furthest below the diagonal. Shaded band: ±0.05 CI around y = x.

### 3.3 The general-purpose model encodes program identity

Holding the classifier fixed isolates the mechanism (Figure 6). Using the same gradient-boosted-tree classifier on every encoder, moving from seen to unseen programs the general-purpose ESM2 loses 0.222 AUC (0.882 → 0.660) — a collapse consistent with embeddings that largely encode which program a sequence belongs to. Under the same classifier, AINN-P1 loses only 0.057 (0.867 → 0.810) and the finetuned ESM2 loses 0.064 (0.880 → 0.816); across all classifiers the median gap is 0.226 for the general-purpose ESM2 versus 0.081 for AINN-P1 and 0.123 for the finetuned ESM2. A general-purpose model trained on all of protein space has no incentive to discard the coarse, easily separable signal that distinguishes one antibody program from another; AINN-P1, built for the antibody domain, instead captures the finer, program-independent biophysical determinants of expression that survive a change of program.

**Figure 6.**
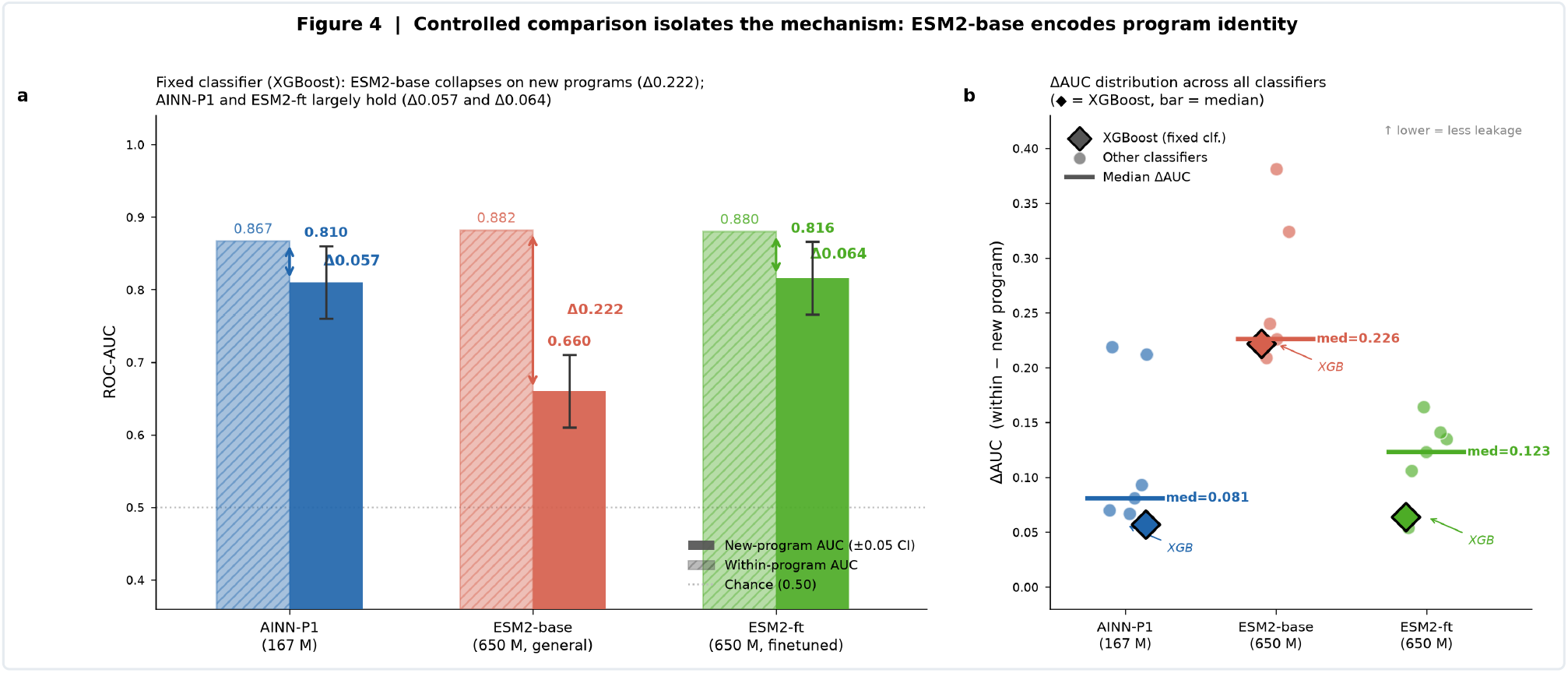
Controlled comparison isolates the mechanism. (A) With a fixed XGBoost classifier the general-purpose ESM2 collapses on unseen programs (Δ0.222, 0.882 → 0.660), while AINN-P1 (Δ0.057) and the finetuned ESM2 (Δ0.064) largely hold. (B) The ΔAUC distribution across all classifiers confirms the effect is not classifier-specific: median gap 0.226 (ESM2-base) versus 0.081 (AINN-P1) and 0.123 (ESM2-ft).

### 3.4 Purpose beats scale for parameter efficiency

Set against parameter count, the efficiency ordering is stark (Figure 7). AINN-P1 delivers its 0.81 new-program AUC at 167 M parameters, exceeding the general-purpose ESM2’s 0.68 at 650 M and matching the finetuned ESM2’s 0.83 at 650 M. On a new-program-AUC-per-parameter basis, AINN-P1 is the clear leader and the general-purpose ESM2 the clear laggard despite being the largest model with the most pretraining data; AINN-P1’s worst tree-model configuration on new programs still exceeds the general-purpose ESM2’s best.

**Figure 7.**
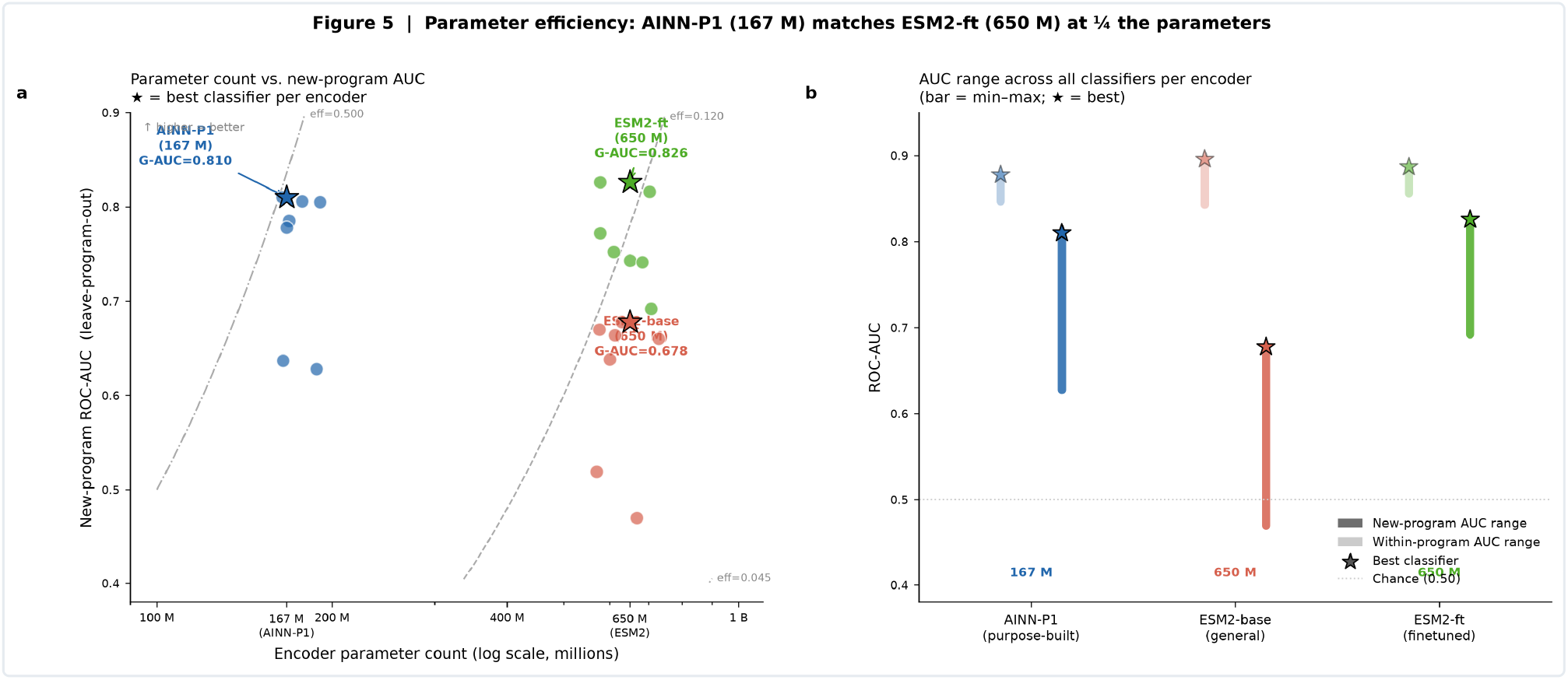
Parameter efficiency. (A) Encoder parameter count (log scale) versus best new-program ROC-AUC; AINN-P1 (167 M) exceeds the general-purpose ESM2 (650 M) and rivals the finetuned ESM2 at far fewer parameters (★ = best classifier per encoder). (B) AUC range across all classifiers per encoder: AINN-P1’s worst tree model on new programs still exceeds the general-purpose ESM2’s best.

**Table 2.** All 21 encoder–classifier configurations plus the majority baseline under both protocols, sorted by new-program AUC. R = random split (within-program); G = grouped, leave-program-out (new-program). Encoders: AINN-P1 (167 M), ESM2-base (general, 650 M), ESM2-ft (finetuned, 650 M). Gap = R-AUC ™ G-AUC.

| Encoder | Classifier | R-AUC | R-Acc | R-MCC | G-AUC | G-Acc | G-MCC | Gap |
| --- | --- | --- | --- | --- | --- | --- | --- | --- |
| ESM2-ft | XGBoost (tuned) | 0.879 | 0.828 | 0.654 | 0.826 | 0.750 | 0.501 | 0.054 |
| ESM2-ft | XGBoost | 0.880 | 0.804 | 0.606 | 0.816 | 0.760 | 0.529 | 0.064 |
| AINN-P1 | XGBoost | 0.867 | 0.794 | 0.584 | 0.810 | 0.706 | 0.404 | 0.057 |
| AINN-P1 | Ensemble (RF+XGB) | 0.873 | 0.799 | 0.594 | 0.806 | 0.701 | 0.394 | 0.067 |
| AINN-P1 | Random Forest (tuned) | 0.875 | 0.833 | 0.664 | 0.805 | 0.725 | 0.455 | 0.070 |
| AINN-P1 | Random Forest | 0.878 | 0.809 | 0.614 | 0.785 | 0.696 | 0.385 | 0.093 |
| AINN-P1 | XGBoost (tuned) | 0.859 | 0.819 | 0.634 | 0.778 | 0.667 | 0.323 | 0.081 |
| ESM2-ft | Ensemble (RF+XGB) | 0.878 | 0.814 | 0.625 | 0.772 | 0.755 | 0.520 | 0.106 |
| ESM2-ft | Random Forest (tuned) | 0.887 | 0.828 | 0.654 | 0.752 | 0.765 | 0.534 | 0.135 |
| ESM2-ft | Random Forest | 0.883 | 0.819 | 0.634 | 0.743 | 0.770 | 0.554 | 0.141 |
| ESM2-ft | Logistic | 0.864 | 0.799 | 0.595 | 0.741 | 0.701 | 0.398 | 0.123 |
| ESM2-ft | PCA50 + Logistic | 0.856 | 0.804 | 0.608 | 0.692 | 0.662 | 0.357 | 0.164 |
| ESM2-base | Random Forest (tuned) | 0.887 | 0.838 | 0.673 | 0.678 | 0.588 | 0.190 | 0.209 |
| ESM2-base | Random Forest | 0.896 | 0.843 | 0.683 | 0.670 | 0.598 | 0.198 | 0.225 |
| ESM2-base | Ensemble (RF+XGB) | 0.889 | 0.838 | 0.674 | 0.664 | 0.613 | 0.230 | 0.226 |
| ESM2-base | XGBoost | 0.882 | 0.809 | 0.616 | 0.660 | 0.623 | 0.248 | 0.222 |
| ESM2-base | XGBoost (tuned) | 0.878 | 0.843 | 0.684 | 0.638 | 0.593 | 0.184 | 0.240 |
| AINN-P1 | PCA50 + Logistic | 0.849 | 0.809 | 0.615 | 0.637 | 0.642 | 0.278 | 0.212 |
| AINN-P1 | Logistic | 0.847 | 0.814 | 0.625 | 0.628 | 0.642 | 0.278 | 0.219 |
| ESM2-base | PCA50 + Logistic | 0.843 | 0.799 | 0.597 | 0.519 | 0.485 | -0.031 | 0.324 |
| ESM2-base | Logistic | 0.851 | 0.775 | 0.549 | 0.470 | 0.471 | -0.057 | 0.381 |
| (any) | Majority baseline | 0.500 | 0.544 | 0.000 | 0.343 | 0.363 | -0.345 | 0.157 |

## 4. Discussion

### A deployable triage tool grounded in a purpose-built representation

AINN-Express turns a simple, deployable pipeline into a useful expression triage tool: given a set of VHH candidates it ranks them by predicted expression so wet-lab capacity targets the most promising, with a human and experiment in the loop rather than as an automated hard filter. Its generalization to unseen programs rests on the AINN-P1 representation, which — as the benchmark shows — outperforms general-purpose protein language models on this task by a wide margin on new programs, at roughly a quarter of the parameters.

### Purpose beats scale for this domain

A 167 M-parameter foundation model generalizes to unseen antibody programs better than a 650 M general-purpose model (0.81 vs 0.68 new-program AUC) and matches a 650 M domain-finetuned model. For antibody property prediction, representation quality — how well the pretraining objective and corpus align with the target domain — matters more than raw parameter count, with the practical corollary that a smaller purpose-built encoder is cheaper to serve, faster to embed with, and easier to deploy.

### The evaluation protocol is decisive

Random splits reward memorization of program identity and hide the gap that leave-program-out reveals; under a random split the three encoders are indistinguishable (∼0.88 AUC). Any antibody-property benchmark reporting only random-split scores is, in effect, measuring how well the encoder recognizes clonal relatives — not how it will perform on the next discovery program. We recommend grouped, leave-program-out (or leave-cluster-out) cross-validation as the default for property-prediction studies on clustered biological panels; the within-program score remains useful as an upper bound and leakage diagnostic, but should never be the headline number.

### Limitations

As a case study, the panel is modest (∼200 sequences), so leave-program-out AUCs carry a confidence interval on the order of ±0.05; AINN-P1’s advantage over the general-purpose ESM2 (Δ0.13) is well outside this band and should be read as real, while its gap to the finetuned ESM2 (Δ0.016) is within the band and should be read as parity. The study covers one antibody format (VHH single-domain) and one property (binary expression), and a small number of sequences carry inconsistent experimental labels, imposing an intrinsic ceiling on accuracy. Predictions should inform prioritization, not replace experimental confirmation.

## 5. Materials and Methods

### 5.1 Model design and training

AINN-Express has two stages. A VHH sequence is first passed through AINN-P1 operating as a frozen encoder, producing one fixed-dimensional embedding with no gradients flowing into the foundation model. A compact gradient-boosted-tree classifier (XGBoost; Chen & Guestrin, 2016) then maps that embedding to an expression probability. Freezing the encoder makes the model fast, reproducible, and data-efficient. The classifier is trained on the available labeled VHH panel with class imbalance handled during fitting; once embeddings are computed the model trains in seconds on CPU, so it is inexpensive to refresh as new expression data arrives.

### 5.2 Encoders (benchmark)

Each sequence is encoded independently by a frozen protein model into a fixed-length vector, with no gradients propagating into the encoder. Three encoders are compared under identical downstream treatment: **AINN-P1** (Ainnocence’s purpose-built protein foundation model, 167 M); **ESM2-base** (a general-purpose ESM2, 650 M, pretrained on UniRef-scale data; Lin et al., 2023; Suzek et al., 2015); and **ESM2-ft** (the same 650 M architecture, domain-finetuned on antibody sequences).

### 5.3 Downstream classifiers

On top of each encoder’s embeddings we fit the same family of seven lightweight classifiers: logistic regression; PCA (50 components) followed by logistic regression; random forests, with and without nested hyperparameter search; gradient-boosted trees, with and without nested search; and a soft-voting ensemble of the random forest and XGBoost models. The same classifier family, grids, and fitting procedure are applied identically to all encoders, so any difference is attributable to the embeddings.

### 5.4 Data and task

We predict binary experimental expression (expressed versus not) for a proprietary panel of approximately 200 VHH sequences spanning multiple antibody-discovery programs; classes are mildly imbalanced (about 54% positive). Candidates within a program are related by design (clonal variants, affinity-matured lineages, framework variants) and are therefore more similar to one another than to candidates from other programs — the clustered structure underlying the leakage analysis. Sequence identities and program details are proprietary and are omitted.

### 5.5 Cross-validation and evaluation

Because candidates within a program are related, a random split can leak near-identical sequences across train and test; we therefore also evaluate leave-program-out, grouping by program so no program is split. Exact-duplicate sequences are collapsed before splitting; hyperparameters are chosen by nested cross-validation so “tuned” scores are not optimistically biased; and metrics are pooled over out-of-fold predictions. ROC-AUC is the primary metric, with accuracy, F1, and MCC reported at a 0.5 cutoff. The AUC gap (random minus grouped) serves as a per-configuration leakage diagnostic. Given the panel size, we estimate a ±0.05 confidence interval on leave-program-out AUCs; differences larger than this band are treated as real, and differences within it as parity.

## 6. Conclusion

AINN-Express predicts VHH expression from sequence alone, reaching ROC-AUC 0.87 within known programs and 0.81 on unseen programs — useful for candidate triage — using a frozen AINN-P1 encoder and a lightweight classifier that needs no per-task training. Its generalization is not incidental: a leakage-aware benchmark shows AINN-P1 embeddings transfer to new antibody programs far better than a larger general-purpose ESM2 and match a domain-finetuned ESM2 at a quarter of the parameters, because purpose-built representations capture transferable determinants of expression where general-purpose ones encode program identity. For narrow biological domains, representation quality can outweigh model scale — provided evaluation is leakage-aware.

## Declarations

### Author contributions

R.W. and K.J.: methodology, software, formal analysis, and writing — original draft. L.P.: conceptualization, supervision, and writing — review and editing.

### Data and code availability

The AINN-P1 model is accessible through Ainnocence’s API. The antibody expression dataset is proprietary and is not publicly released.

## Acknowledgments

The authors thank the Ainnocence research and engineering teams.

